# Augmentation of T-cell activation by oscillatory forces and engineered antigen-presenting cells

**DOI:** 10.1101/580704

**Authors:** Fatemeh S. Majedi, Mohammad Mahdi Hasani-Sadrabadi, Timothy J. Thauland, Song Li, Louis-S. Bouchard, Manish J. Butte

## Abstract

Activation of T cells by antigen presenting cells allows them to proliferate, produce cytokines, and kill infected or cancerous cells. We and others have shown that T cell receptors receive and in fact require mechanical forces from their own movements and the movements of antigen presenting cells. Emulation of T cell activation in vitro allows for the massive expansion of T cells necessary for clinical applications. In this paper, we studied the impact of augmenting novel artificial antigen presenting cells of various sizes and antigenic signal strength with mechanical, oscillatory movement. We showed that dynamic culture roughly doubles signal strength as compared to conventional, static culture. We demonstrated that tuning the strength of signal to a “sweet spot” allows for robust expansion of induced regulatory T cells, which is impeded by approaches that simply maximize activation.

## Introduction

T lymphocytes circulate throughout the body and coordinate the immune response against pathogens by recognizing their proteome as foreign. Activation of T cells begins by T cell receptors (TCRs) engaging with antigenic peptides proffered by the major histocompatibility complex (p-MHC) of antigen presenting cells (APC). Triggering of the TCR by pMHC requires a mechanical force that pulls upon the TCR (Liu et al., 2014; Kim et al., 2009). Those pulling forces are at least partially made by the T cells themselves through a series of oscillatory pulling movements (Hu and Butte, 2016). We have previously shown that these forces can be substituted by an exogenous source, for example, ligands tethered to the cantilever of an Atomic Force Microscope (Hu and Butte, 2016). Those experiments entailed contact with a single T cell at a time, and thus did not allow for scaling the application of forces toward the large numbers of T cells needed for industrial, clinical purposes. In this paper, we investigated if provision of exogenous forces *in vitro* could augment the force-based triggering of TCRs in a large number of T cells.

*Ex vivo* cultivation of T cells is important for manufacturing cellular therapies, such as CAR-T cells, so optimizing approaches for polyclonal T cell cultivation are clinically important (Xu et al., 2018). To activate T cells for engineering, they are commonly cultured with beads coated with stimulatory antibodies, or artificial antigen presenting cells (aAPCs). A number of groups have developed aAPCs using micro- and nanotechnology approaches, developing particles that can be co-cultured with T cells or engineered surfaces that offer stimulatory signals (Perica et al., 2015). In this work, we compared our particles with CTS Dynabeads (TM), which are the most popular, commercial aAPC, roughly the size of resting T cells (4-5 μm diameter), comprising a rigid, polystyrene shell around a superparamagnetic particle, and coated with stimulatory antibodies.

The key to T-cell activation is offering simulation to the TCR, either in the form of pMHC or by using antibodies that trigger the CD3 chains of the TCR complex. Naïve T cells require secondary stimulation of the CD28 receptor for complete activation, and so antibodies that cross-link and activate CD28 are almost always included in the formulation of aAPCs. The amount of signal provided by the aAPC is proportionate to their cost, and so many approaches have attempted to identify and minimize the amount of signal needed. T cells stimulated on a nanopattered antigen array comprising a planar lipid bilayer showed that the number of antigens on a surface is more important than the density, with a density of 90–140 pMHC/μm^2^ yielding maximal stimulation (Deeg et al., 2013). Nanoarrays formed by block copolymer micellar nanolithography (BCML) functionalized with gold nanoparticles found a plateau of maximal response when the distance between stimulatory signals was 60 nm or less (Matic et al., 2013). Surface curvature, however, may contribute to T cell activation (He and Bongrand, 2012; Rossy et al., 2018), and so it is difficult to extrapolate antigen amounts from experiments on planar surfaces.

Here, we manufactured spherical aAPCs of different sizes to offer various degrees of curvature. We conjugated these particles with anti-CD3 and anti-CD28 at different densities to measure the effect of strength of stimulatory signals on T-cell activation. We also delivered an oscillatory stimulus to the culture to test whether an external mechanical stimulus upon the TCR can promote activation. Overall, we found conditions that dramatically improved activation beyond conventional, Dynabead-based stimulation. In another example, we chose aAPC conditions that offered a “sweet spot” of signaling to maximize the production of induced regulatory T cells (iTreg), the development of which is actually hindered by high levels of stimulation. The particles were also endowed with the ability to secrete cytokines to further promote iTreg development. These examples show that delivery of mechanical forces coupled with aAPCs of tunable size/curvature, signal density, and cytokine section offer the possibility to fruitfully engineer T cells for a variety of clinical and experimental needs.

### Design considerations

An important design consideration was endowing the particles with the ability to be easily separated from cells during modification, washing steps, and especially after co-culture. Thus, we made the beads magnetically responsive by embedding superparamagnetic iron oxide nanoparticles (SPIONs) during their microfluidic synthesis. The processing conditions were tuned to encapsulate around 2.8 ± 0.3 Vol% SPIONs per particle (**Table SI.1**), which was identified in pilot work to be sufficient for magnetic separation.

Our polymeric, microparticle aAPCs are composed of alginate, which is ionically crosslinked by addition of divalent cations like calcium. Ionic crosslinking of alginate provides sufficient working stability; however, we saw irreversible changes (hysteresis) in particle morphology during pilot magnetic separation and traction experiments. To overcome this issue, we used our recently developed combinatorial crosslinking technique which provided both chemical and physical crosslinking (Hasani-Sadrabadi et al., 2018).

## Results and Discussion

To develop our artificial APCs, we utilized a microfluidic droplet generator that encapsulates alginate polymer and magnetic nanoparticles (**Fig. 1A**). A constant fraction of crosslinker (4-arm PEG hydrazide) was mixed with the alginate polymer in the main channel. During droplet formation, we encapsulated within our beads magnetic nanoparticles that were 100 nm-diameter, carboxylated superparamagnetic iron oxide nanoparticles (SPIONs). The microfluidic approach produced a homogeneous collection of particles, as verified by transmission electron microscopy (TEM) (data not shown) and dynamic light scattering (**Fig. 1C**). To tune the size of particles ranging from 150 nm to 10 μm, we varied the ratio of central to sheath flow (**Fig. 1B**). To cover a broad range of particle sizes (over one order of magnitude), we selected three sizes of particles with average radii of 307 nm (“0.3”), 824 nm (“0.8”), and 4540 nm (“5 μm”) (**Fig. 1C**). In pilot work, we found that nanoparticles of 50-250 nm diameter could not be fully separated from one another or from T cells due to partial internalization or trapping on the rough cell surface, whereas particles larger than 300 nm could be separated from cells with >95% efficiency (not shown). The resulting alginate particles were then collected in a bath of 200 mM CaCl_2_, followed by a ~40 min incubation to reach complete gelation. Eventually particles were subjected to overnight chemical crosslinking though the hydrazine linker. Excess calcium and crosslinker were removed by serial washing with phosphate buffered saline.

**Figure 1.**
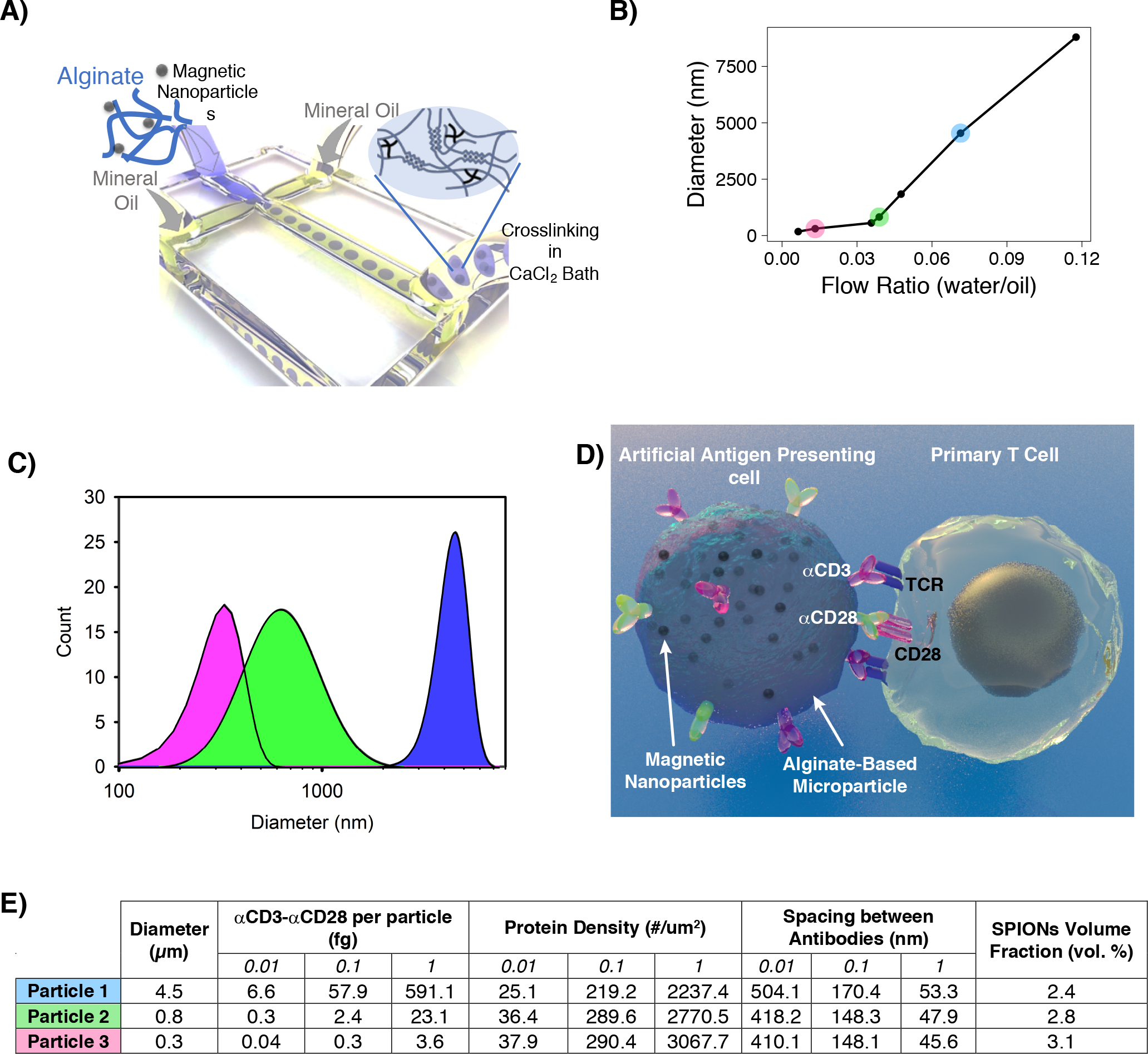
(A) Schematic representation of microfluidic generation of alginate nano-/microparticles encapsulating magnetic nanoparticles. (B) Size distribution analysis of prepared microparticles at different flow rates. (C) Size distribution analysis of selected particles (0.3, 0.8, 4.5 *μ*m). (D) Schematic representation of proposed interactions between artificial antigen presenting cells and primary T cells. (E) Summary of physical characteristics of prepared library of particles to present broad range of antigens on their surfaces.

We next conjugated stimulatory antibodies to the surface of the microparticles. The carboxylic groups of alginate provide a versatile platform for antibody conjugation. Using NHS/EDC chemistry, we conjugated anti-CD3 and anti-CD28 antibodies and washed away excess antibodies and quenching unreacted groups through repeated washing with phosphate buffered saline containing 0.5% BSA. Based on pilot experiments, we chose three different densities of antibodies to coat beads representing low, medium, and high amounts of antigenic signaling. An average of 2692 ± 420, 266 ± 41, and 33 ± 7 antibody molecules per square micrometer were immobilized as high, medium, and low conjugation densities, respectively (**Fig. 1E**). For comparison, a theoretical limit of ~12,732 antibodies could be packed antibodies into a square micrometer, assuming the antibody has a ~5 nm radius (Reth, 2013), so the “high” labeling density here is approximately 21% of the limit. The size of the interface between T cells and antigen presenting cells (**Fig. 1D**) varies based on cytoskeletal state of the T cell (Thauland et al., 2017), with most contacts falling in the range from 5-25 μm^2^. If one assumes an area of 10 μm^2^ for a typical immune synapse, the large particles (2.25 μm radius) would offer a hemispheric area of ~32 μm, so an immune synapse sized 10 μm^2^ would engage ~⅓ of the hemisphere and would engage ~26K antibodies (high density coating), 2660 antibodies (medium), and 330 molecules (low). For the medium-sized particles (0.4 μm radius) a hemispheric immune synapse offers ~1 μm^2^ area and 2692, 266, and 33 molecules, at the respective conjugation densities. For the smallest particles (0.15 μm radius), a hemispheric immune synapse would engage 376, 37, and 5 molecules, respectively. Thus, in all cases, the number of engaged T cell receptors is greater than the experimentally observed minimum amount of signal needed to activate a T cell, which is a single engage TCR (Irvine et al., 2002).

To assess the impact of external, gentle mechanical stimulation on co-cultures of T cells with aAPCs we used an orbital shaker to deliver a continuous oscillatory movement of either ~250 rpm rotational speed (“dynamic”) or switched off (“static”) (**Fig. 2A**). The impact of the aAPCs on T cells were compared with that of the popular, commercially available CD3/CD28 T-cell expansion beads (Dynabeads from Life Technologies). Primary T cells were obtained from spleens of wild-type mice, enriched by negative magnetic-bead selection, and cultured with aAPCs in either static or dynamic conditions. During our experiments we kept the particle dose appropriate to provide a constant activation surface area of 9.5×10^7^ *μ*m^2^ per 1.5×10^6^ T cells (**Table SI.2**). This calculation was equivalent to the surface area of Dynabeads in co-culturing one Dynabead per one primary T cell, the manufacturer’s recommendation.

**Figure 2.**
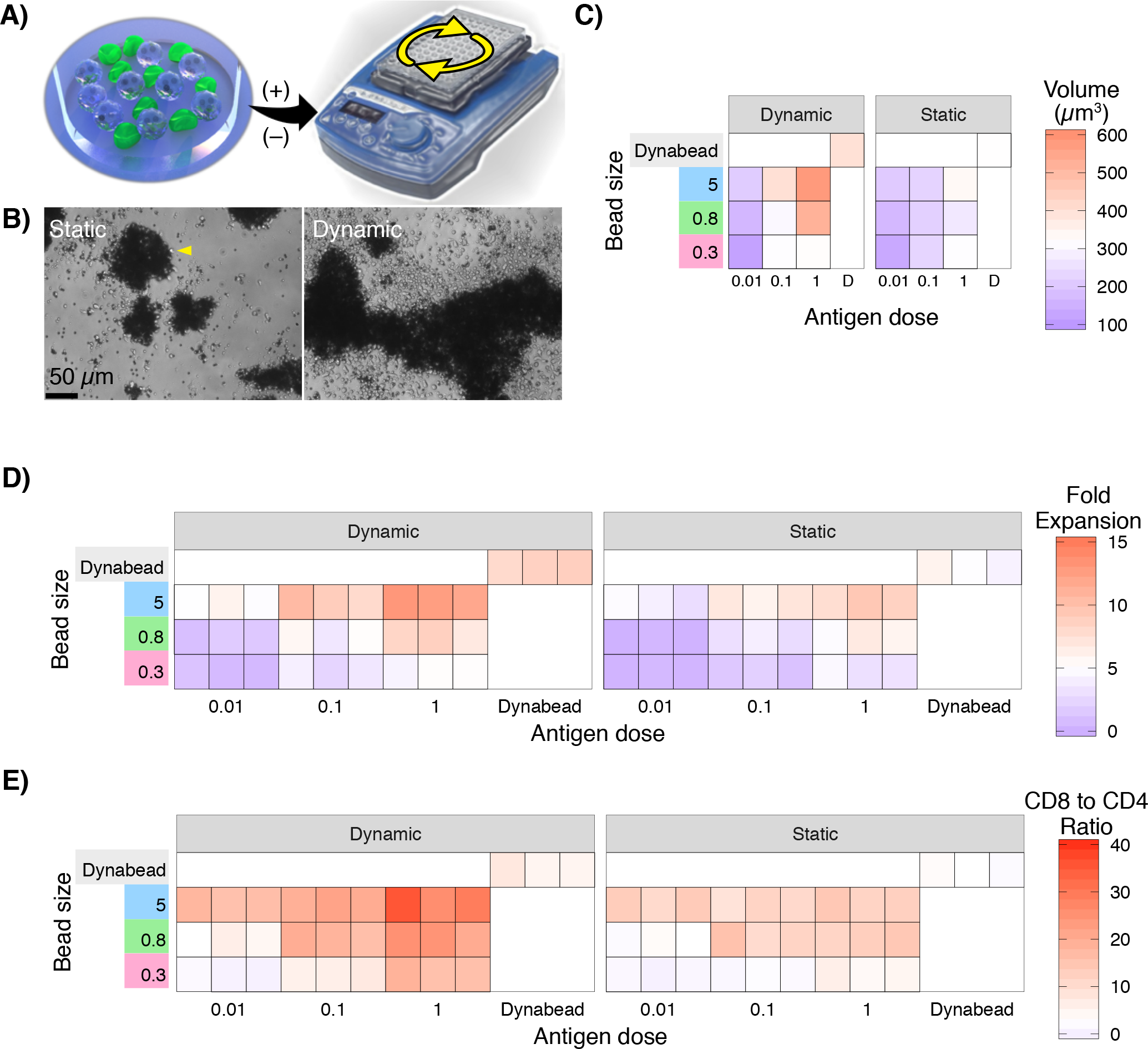
(**A**) Particles and T cells were co-cultured under static and dynamic conditions. (**B**) Representative brightfield microscopy images of formed clusters by primary mouse T cells cultured with 5 μm aAPCs at a constant dose (1:1 particle/ T cell ratio) under static and dynamic (shaking) cultures. Scale bars, 50 μm. (**C**) The mean volumes of T cells activated and expanded using various formulations of particles, as indicated. (**D**) Expansion of primary mouse CD4+ T cells by varying the antigen dose, particle size or the culture conditions after four days. (**E**) FACS quantification of CD4/CD8 ratio of CD4^+^ and CD8^+^ single-positive cells cultured with varying formulations of particles compared to Dynabeads.

By day 2-3 of culture, polyclonal, primary mouse T cells formed large clusters of beads and cells. The clusters were obviously larger in the dynamic culture than in static culture (**Fig. 2B**). The T cells were separated from the beads and imaged by 3D confocal microscopy to assess their growth. Cell volume was assessed because volume changes as a cell grows, proliferates, or differentiates. Cell volume can also change in response to external physical cues (Guo et al., 2017). We found that cell growth was larger in dynamic culture versus static culture across all particle sizes and conjugation densities (**Fig. 2C**). The average volume of T cells co-cultured with Dynabeads in static conditions (n=22) was 321 μm^3^, which represents an average diameter of 8.5 +/− 0.34 μm. The largest cells were those cultured with 4.5 μm particles with the highest density of ligands, an average diameter of 10.3 +/− 0.83 μm (n=25).

To assess their proliferative response, we counted T cells after 3 days of co-culture with the various particles. In all conditions, dynamic culture resulted in significantly higher expansion of T cells than static culture (**Fig. 2D**). The average fold expansion of T cells co-cultured with Dynabeads in static conditions (n=3) was 5 +/− 1.8 (mean +/− 95% CI) fold. The largest fold expansion was observed in the condition where T cells were cultured with our 4.5 μm particles with the highest density of stimulatory antibodies, resulting in an increase of 12.5 +/− 1.2 fold. Averaging across all particle sizes and antigen doses, shaking increased the proliferation of the cells by 2.0- fold compared to static culture (ANOVA considering movement, size, and dose; movement p = 1.5 × 10^−12^).

In general, cytotoxic CD8+ T cells have a higher proliferative capacity than CD4+ T cells. Cytotoxic T cells have important applications in engineered cancer immunotherapies. We assessed the ability of these particles to promote cytotoxic T cell expansion by monitoring the CD8-to-CD4 T cell ratio during proliferation. We separately purified CD4+ T cells and CD8+ T cells from mice, then mixed them to exactly to achieve the physiological ratio of one CD8+ T cell to two CD4+ T cells. We co-cultured T cells with particles as above, and after 5 days, we measured the ratio of CD8 to CD4 T cells by flow cytometry. The average CD8-to-CD4 ratio of T cells co-cultured with Dynabeads in static conditions (n=3) was 2.75 +/− 1.5 (mean +/− 95% CI). The largest increase in the cellular ratio was observed in the condition where T cells were cultured with 4.5 μm particles with the highest density of ligands, resulting in a CD8 to CD4 ratio of 30.1 +/− 9.8. Averaging across all particle sizes and antigen doses, shaking increased the CD8 to CD4 ratio of the cells by 2.1- fold compared to static culture (ANOVA considering movement, size, and dose; movement p < 10^−16^).

We noted that the larger particles resulted in more expansion, and especially CD8 expansion, of the T cells than the smaller particles, even though the density of antibodies across the beads of different sizes was almost identical (**Fig. 1E**) (ANOVA size p < 2 × 10^−16^). This result shows that the immune synapse integrates the aggregate number of molecular signals across the interface, rather than the density of antigenic ligands.

We further examine the proliferative responses of T cells in response to stimulation with our aAPCs, we used a dye-dilution approach to follow the proliferation pattern. Sequential generations of daughter cells result in roughly two-fold dilution of the fluorescent signal (**Fig. 3A**). The average %proliferation of T cells co-cultured with Dynabeads in static conditions (n=3) was 91.0 +/− 5.8% (mean +/− 95% CI) (**Fig. 3B**). The maximum proliferation was observed in the condition where T cells were cultured with 4.5 μm particles with the highest density of antibodies, resulting in proliferation of 98.8 ± 1.9%. Averaging across all particle sizes and antigen doses, shaking increased the %proliferated of the T cells by 1.72- fold compared to static culture (ANOVA considering movement, size, and dose; movement p = 0.011). We also examined expression of T-cell activation markers CD25 and CD44 by flow cytometry after activation as well. We found that expression of these activation markers trended similarly to proliferation (**Fig. 3D-G**). As with absolute expansion, activation and proliferation were greater for larger beads than smaller beads even when antibody density is held constant. Together these results show that activation and proliferation are proportional to the amount of antigen rather than its density.

**Figure 3.**
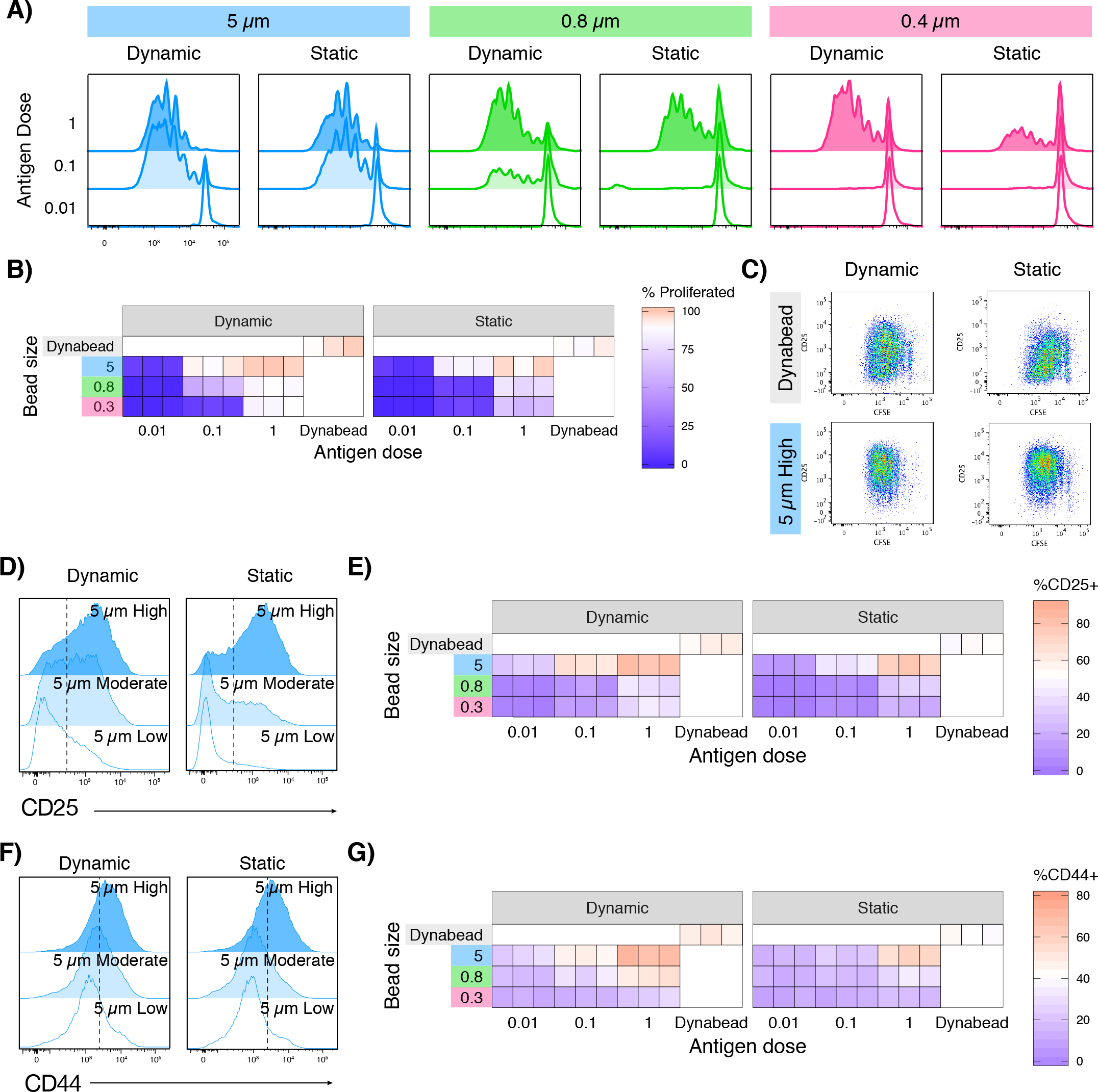
Proliferation and activation analyses of CD4^+^ T cells cultured under static or dynamic conditions in the presence of varying formulation of particles. (**A**) Flow cytometry histograms CFSE dilution and (**B**) Percentage of proliferated T cells three days after co-culturing with different formulation of engineered particles. (**C**) (**D**) CD25 expression histograms after 24 h of co-culturing of primary naïve CD4+ T cells with 5 μm microparticles presenting various surface density of antibodies under static or dynamic culture. (**E**) Percentage of CD25+ T cells 24 h after activation with various formulation of particles or Dynabeads. (**F**) CD44 expression histograms after 24 h of co-culturing of primary naïve CD4+ T cells with 5 um microparticles presenting various surface density of antibodies under static or dynamic culture. (**G**) Percentage of CD44+ T cells 24 h after activation with various formulation of particles or Dynabeads.

We showed in prior work that the size of the immune synapse in naive versus effector (recently activated) T cells controls the amount of signal they accumulate in their interactions with APCs (Thauland et al., 2017). We sought to test the coupling of signal accumulation and synapse size in a reductionist manner. After OT-II T cells were activated with particles, as above, for 24 hours, we purified away the stimulatory particles and co-cultured them with the B cell lymphoma line LB27.4 that was loaded with ovalbumin peptide antigen. We measured the average volume of immune synapses formed between T cells activated in different culture conditions and B cells. based on the accumulation of the integrin leukocyte function–associated antigen 1 (LFA-1) measured by the volume of positive pixels (**Fig. 4A**). The average synapse size for T cells co-cultured with the 4.5 μm particles at high levels of stimulatory antibodies in static conditions (n=3) was 32.8 ± 4.1 μm^2^ (**Fig. 4B**). The maximum proliferation was observed in the condition where T cells were cultured with 4.5 μm particles with the highest density of antibodies in shaking conditions (n=3) 36.7 ± 4 μm^2^. Averaging across all particle sizes and antigen doses, shaking increased the percentage of proliferated T cells by 1.72- fold compared to static culture (ANOVA considering movement, size, and dose; movement p = 0.011). These results show that the size of the immune synapse is larger when the cells are more activated. Our prior published result compared the size of the synapses for naïve T cells versus effector T cells (lymphoblasts in day 3-5 of culture) (Thauland et al., 2017), and showed that activation of cofilin in effector T cells enabled changes to the cytoskeleton that allowed for larger synapses and a lower threshold of activation than naïve T cells. In other words, that the threshold of activation was determined by the size of the synapse and thus the dynamic ability of the cytoskeleton to rearrange when in contact with an APC. The result here goes further: we demonstrate here that the dynamic size of the synapse is not just binary, but rather gradated based on signal strength.

**Figure 4.**
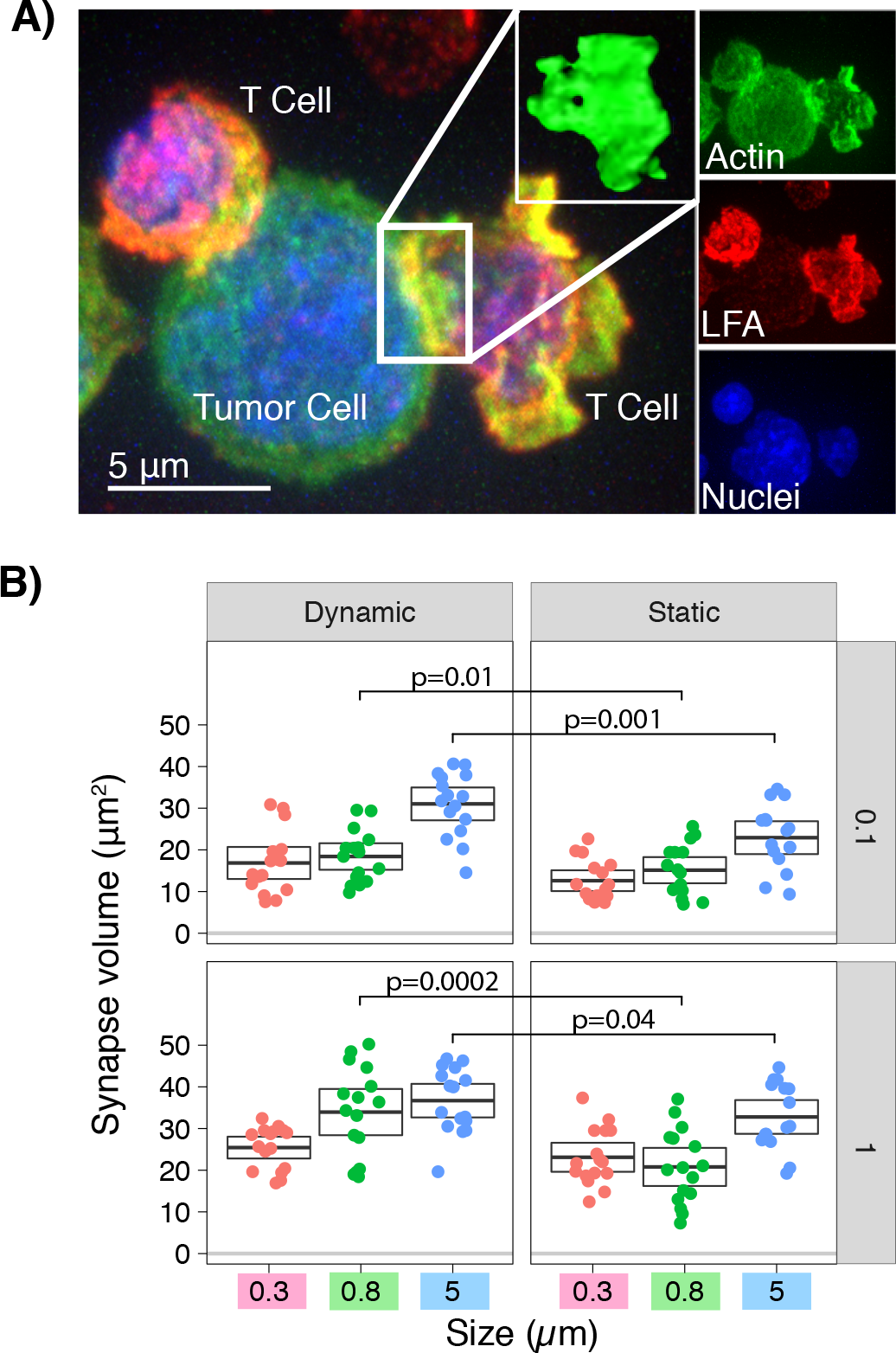
(**A**) Immune synapses formed by OT-II T cells activated with 5 *μ*m (1) microparticles interacting with (antigen-pulsed) antigen presenting cells (B lymphoma cells) were imaged by confocal microscopy. Images show overlap of confocal slices. Representative cells that had the median immune synapse volume were chosen. (**B**) Analysis of immune synapse volumes (in *μ*m^3^) formed by primary naïve T cells activated with various particle sizes with high or low antibody conjugation level cultured at static or dynamic conditions. Each dot represents an immune synapse between a T cell and an antigen presenting cell. Boxes show means and 95% CI values. Results are representative of three independent experiments.

Activating T cells with high signal strength allows for massive expansion, which is needed for transducing chimeric antigen receptors (CARs) and having sufficient transduced cells for a therapeutic dose. The opposite problem arises when expanding T cells in culture for the purposes of generating engineered regulatory T cells. In vivo, regulatory T cells can be elicited to foreign antigens when they are provided at low levels, rather than at high signal strength (Rubtsov et al., 2010). Induction of regulatory T cells is improved by provision of TGF-β and IL-2 (Zheng et al., 2014). We recently demonstrated that alginate microparticles could be loaded with cytokines to skew T cells to iTregs (Majedi et al., 2018), which served as a motivation to combine that capability with the tunable signal strength of the system presented here. To evaluate the effect of mechanical forces and antigen strength on iTreg formation, we loaded Alg-Hep particles with TGF-β and IL-2. We sorted rigorously naïve CD4+ T cells to eliminate natural regulatory T cells, and then co-cultured these cells with our microparticles. Dynabeads were used as the comparison, providing an equivalent amount of soluble TGF-β in the media.

We assessed the development of Tregs by intracellular staining for the key transcription factor Foxp3 followed by flow cytometry. The mean fluorescence intensity of the Foxp3 transcript correlates to their regulatory ability (Chauhan et al., 2014), and so Foxp3 expression level was measured as well.

The percentage of iTregs induced in culture with Dynabeads plus TGF-β in static conditions (n=3) was 25.7 +/− 10.1% (mean +/− 95% CI) (**Fig. 5A**). The maximum iTreg induction was observed in the condition where T cells were cultured with 4.5 μm particles with the lowest density of antibodies, resulting in iTreg induction of 71.6 ± 9.6%, almost three-fold higher. In general, averaging across all particle sizes and antigen doses, shaking did not alter the rate of iTreg induction at all (1.02-fold difference). Curiously, the very lowest amounts of signal strength, as seen in proliferation and activation assays above, were not able to elicit high yields of iTreg induction. In fact, a “sweet spot” of signal was needed, either in the form of larger beads with lower antibody coating or smaller beads with higher antibody coating amounts (**Fig. 5A**). The expression of Foxp3 on a per-cell basis was highest in the conditions that elicited the highest induction of iTregs (**Fig. 5B**).

**Figure 5.**
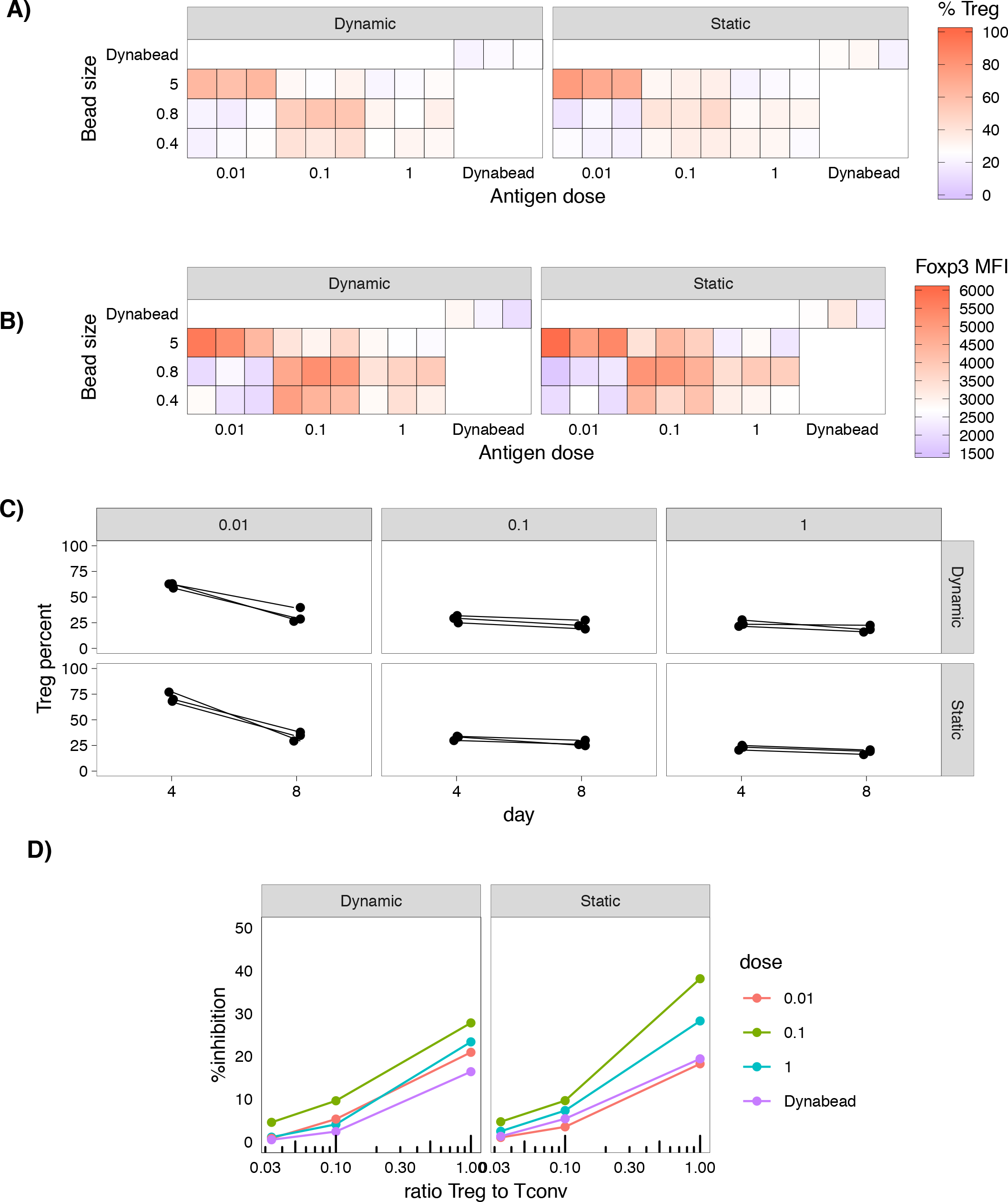
(**A**) Release profile of TGF-β from alginate-based particles at 37 °C. TGF-β releasing particles generate a sustained signal to induce formation of induced regulatory T cells (iTreg cells). Flow cytometric analysis of iTreg development was assessed by flow cytometry for Foxp3 and CD25 co-expression after co-culture of naïve CD4+ T-cells with particles at various formulations either in dynamic or static conditions for 4 d. (**A**) Percentage of induced Tregs and (**B**) mean fluorescence intensity (MFI) of Foxp3 expression in T cells 4 d after activation with various formulation of particles or Dynabeads. (**C**) Stability of formed T-regs as assessed by measuring the change in the population of iTregs (T cells expressing CD4, CD25, and Foxp3+) after 4 and 8 days. (**D**) T cell suppression assay. Flow sorted Tregs co-cultured with naïve primary CD4+ T-cells (T-conv) at three different ratios of (1:1, 1:10, and 1:30 of T-reg to T-conv) in the presence of surface coated anti-CD3 and soluble anti-CD28 for 3 d.

We examined the stability of the Foxp3 protein expression, because transient expression of Foxp3 does not yield highly suppressive regulatory T cells (Rubtsov et al., 2010). The T cells generated through culture with the 4.5 μm particles were separated from the particles and then maintained in culture with IL-2. We assessed for expression of Foxp3 by flow cytometry at day 4 and at day 8 of culture (**Fig. 5C**). We found that expression of Foxp3 was dramatically reduced in the condition where the induction was highest, by the 4.5 μm particles that offered the lowest antigen signal (0.01). Dynamic culture mitigated the loss of Foxp3 expression modestly (29.9 ± 13.1% decrease in dynamic culture versus 37.6 ± 17.7% in static culture). In contrast the antigenic strengths that were moderate (0.1) and highest (1) had the most stability in culture (4-5% diminshment). Overall, at day 8, the highest expression of Foxp3 was still seen in the co-culture conditions with 4.5 μm particles that offered the lowest antigen signal (0.01), 34 ± 8.8% (n=3).

The ultimate in vitro test of regulatory T cell function is assessed by their ability to suppress the effector responses of conventional, activated T cells. We co-cultured iTregs induced in a variety of conditions above with conventional T cells at a cellular ratio of 0, 1, 10 and 30 CFSE-labeled conventional, naïve T cells to one iTreg and stimulated with anti-CD3 and anti-CD28. Proliferation of the naïve T cells was assessed without Tregs, and % inhibition was measured by subtracting the proliferation as seen when Tregs were co-cultured. We found maximal inhibition, and thus maximal regulatory function, was enacted by the iTregs that were generated in the “sweet spot” condition of culture with 4.5 μm particles with middle levels of activating antibodies (0.1) (**Fig. 5D**). Together these results show that activation and generation of regulatory T cells can be optimized by culturing with particles of large size but low to middle antigenic strength, resulting in the highest stability of regulatory T cells and the most inhibition of effector T cell proliferation.

## Conclusion

T cells form an interface with antigen presenting cells (APCs) called the “immune synapse” that allows for triggering of TCRs by pMHCs. Artificial antigen presenting cells (aAPCs) can be fabricated to emulate this response and allow for productive expansion of T cells in vitro. By manipulating the size and density of stimulatory signals on aAPCs, we can learn about what is required in the TCR-pMHC interface to reach the threshold for T cell activation and thus improve cultivation of engineered T cell therapies. No other work, to our knowledge, has explored the combination of artificial antigen presenting cells with mechanical stimulation as we did here. We showed overall that gentle mechanical stimulation offers an approximately 2-fold increase in signal strength to T cells as compared to conventional static cultures.

Other work in this area has demonstrated the ability to maximize signaling and proliferation, akin to the work presented in the first part of this paper. AAPCs fabricated from biomimetic scaffolds, namely mesoporous silica nanorods coated with a lipid bilayers, allowed for maximizing signal strength and an allowed for impressive expansion of T cells *in vitro* (Cheung et al., 2018), but did not allow for obviously tuning down the signal as we did here. Another aAPC system was demonstrated recently with the conjugation of pMHC onto the surface of yeast cells, and this work revealed that density of pMHC was correlated to activation of T cells (Smith et al., 2018). Conversely, our work here and other reductionist approaches (Manz et al., 2011) demonstrated that absolute amount of antigenic signal, rather than density, is more important. Nano-aAPCs of 100 nm diameter coated with anti-CD3 and CD28 were able to elicit modest proliferation *in vitro* but in the *in vivo* context could traffic down lymphatics to draining lymph nodes more readily than micron scaled particles (Rhodes and Green, 2018). Recent work with aAPCs comprising polyisocyanopeptide polymers 100-1000 nm long, functionalized with anti-CD3 and IL-2, showed that closer spacing of the anti-CD3 and IL-2 contributed to T-cell activation (Hammink et al., 2018). These aAPCs together reveal that tuning antigenic signals and spatial separations has utility and offers an advantage over biological antigen presenting cells when developing engineering solutions to scaling the production of T cells for therapeutic purposes.

Others have shown that the mechanical stiffness of the surface where antigens or antibodies are tethered has a significant impact on the proliferation and effector responses of T cells. In response to polyacrylamide (PA) surfaces of different stiffness coated with anti-CD3 antibodies, T cells appear to proliferate strongly in the context of stiff PA surfaces rather than soft ones (Saitakis et al., 2017). Our work utilized mechanically soft particles (alginate) as well as extremely stiff, commercially available ones (Dynabeads) and found that culture with softer particles could actually outperform stiffer ones, though the stiffness axes was not directly probed in our work and could be explored in future work as alginate is quite tunable for mechanical stiffness (Majedi et al., 2018). The ability of conventional Dynabeads or stiff microparticles to serve as a source of cytokines is inherently limited, as shown by recent work demonstrating that IL-2 activity is abrogated when immobilized on very rigid aAPCs (Hammink et al., 2018). Thus, there may be a net advantage in utilizing the cytokine-secreting capability of relatively soft alginate particles plus their ability to elute cytokines.

## Supporting Information

### Materials and Methods

#### Experimental Procedure

##### 1. Chemicals and Biologicals

Unless noted otherwise, all chemicals were purchased from Sigma-Aldrich, Inc. (St. Louis, MO). All glassware was cleaned overnight using concentrated sulfuric acid and then thoroughly rinsed with Milli-Q water. All the other cell culture reagents, solutions, and dishes were obtained from Thermo Fisher Scientific (Waltham, MA), except as indicated otherwise.

##### 2. Preparation and characterization of cell-mimicking microparticles

The alginate was charcoal treated, and sterile filtered (0.22 μm filters, Millipore, Billerica, MA) prior to the particle formation. A hydrophobic glass microfluidic droplet junction chip (channel depth 100 *μ*m; Dolomite Microfluidics, Charlestown, MA) was utilized to make monodispersed hydrogel droplets as microparticle substrates. A mixture of alginate solution (1% w/v) and 4-arm PEG hydrazide (MW 5 kDa, Creative PEG work, Chapel Hill, NC) (5 mM) was used as the inner aqueous phase. Mineral oil containing 10 wt% surfactant Span 80 was used as the continuous phase. Several flow rates were applied using two syringe pumps (Fusion 200, Chemyx, Stafford, TX) for the alginate and oil flows to control formation of alginate-based droplets. Images were taken at various time points using a Leica DMIL inverted fluorescence microscope fitted with appropriate filters and connected to a camera. Once particles were formed, they were collected in a bath containing calcium ions (100 mM CaCl_2_) and left at room temperature for 45 min for ionic crosslinking. The microgels were extensively washed with 10 mM NaCl solution and centrifuged (15,000 rpm for 10 min) twice before further incubation in a solution containing hydroxybenzotriazole (HOBt) and 1-ethyl-3-(3-dimethylaminopropyl)carbodi-imide (EDC). After 2 h, particles were dialyzed against deionized water for three days extensively to remove any residual reagents, then frozen at −20 °C and lyophilized. Particles were then resuspended either in deionized water or phosphate-buffered saline (PBS) for further use. Magnetic microparticles were fabricated with the addition of super paramagnetic iron oxide nanoparticles (SPION; 50 nm, carboxylated, Chemicell GmbH, Berlin, Germany) to the alginate/PEG mixture. The solution was (bath) sonicated for 10 min at 4 °C prior to use.

For the preparation of antibody-conjugated alginate particles, EDC/NHS chemistry was used to covalently conjugate anti-CD3 (2C11; Bio X Cell) and anti-CD28 (37.51; Bio X Cell) to the surface of particles. After activation of particles’ carboxylic groups for 10 min and washing them with PBS (1×) twice, these proteins were added to the particles and vortexed briefly before stirring overnight at 4 °C. The protein-functionalized microparticles were then magnetically separated from unbound proteins and washed several times with PBS (1×). Unreacted functional groups were quenched by washing samples in Tris buffer (100 mM, pH 8).

The antibody density that is used for conventional plate-bound stimulation (0.01 μg/mL) was used as the maximum amount of antibody density employed (“1” in Figure 1E). Dilutions of 10- and 100-fold were made as medium and low conjugation densities. The number of particles at different sizes were normalized to their surface area in order to provide the same surface area for protein conjugation and also for co-culturing with T cells. The amount of protein that was used is summarized in the following table:

| | Diameter ($\mu\text{m}$) | Surface area ($\mu\text{m}^2$) | Required total protein per particle (fg) | Number particles per test* | Total aCD 3 ($\mu\text{g}$) | Total aCD 28 ($\mu\text{g}$) | Particle Stock Concentration (#/ml) | Required # of particles |
| --- | --- | --- | --- | --- | --- | --- | --- | --- |
| Particle 1 | 4.5 | 63.62 | 795.19 | $1.5 \times 10^6$ | 1.19 | 0.30 | $1.45 \times 10^8$ | $1.5 \times 10^6$ |
| Particle 2 | 0.8 | 2.01 | 25.13 | $51.9 \times 10^6$ | 1.30 | 0.33 | $248 \times 10^8$ | $1642.2 \times 10^6$ |
| Particle 3 | 0.3 | 0.28 | 3.53 | $193.7 \times 10^6$ | 0.68 | 0.17 | $2150 \times 10^8$ | $43582.1 \times 10^6$ |

\*: per $1.5 \times 10^6$ T cell

Quantification of the total amount of aCD3/aCD28 presented on functionalized particles was analyzed using micro-BCA assay according to the manufacturer’s protocol. Dynamic light scattering (DLS) and zeta potential measurements were performed using a Zetasizer (Zetasizer 3000HS, Malvern Instruments Ltd., Worcestershire, UK) in backscattering mode at 173° for the diluted suspensions in water. The iron content of the particles was measured by ICP-OES (Inductively Coupled Plasma – Optical Emission Spectrometry) after digestion with alginate lyase.

To prepare microparticles loaded with IL-2 and TGF-β, crosslinked particles were incubated with the proteins in PBS buffer containing bovine serum albumin (BSA; 0.1 %w/v) and were gently shaken overnight at 4 °C. The particles were then centrifuged and washed several times to remove unabsorbed proteins. The concentrations of IL-2 and TGF-β in the removed supernatant were measured using enzyme-linked immunosorbent assay (ELISA) to estimate the binding capacity of particles. To study the in vitro release profile, protein-loaded particles were dispersed in PBS (pH 7.4) and 500 μL of particles were placed in Eppendorf tubes, gently shaken, and incubated at 37 °C. At predetermined time points, samples were collected using centrifugation and the supernatant was replaced with an equivalent volume of fresh PBS solution. The concentrations of released proteins from the particles were determined using human IL-2 TGF-β ELISA kits.

##### 3. Co-culture of particles and immune cells

###### 3.1. T cell isolation and activation

Five- to eight-week-old wild-type (C57Bl/6) mice were purchased from the University of California, Los Angeles (UCLA) and maintained in specific pathogen-free facilities at UCLA. All experiments on mice and cells collected from mice were performed in strict accordance with UCLA’s institutional policy on humane and ethical treatment of animals. Cell culture media was RPMI supplemented with 10% heat inactivated FBS, 1% penicillin/streptomycin, 1% sodium pyruvate, 1% HEPES buffer, 0.1% uM 2-ME. Total T cells, CD4+ T cells or CD8+ T cells were purified using negative enrichment kits (Stem Cell Technologies).

Standard Dynabeads or microparticle-assisted in vitro activation of purified T cells (CD4+) were used in this study. Standard in vitro activation of purified T cells (CD8+ or CD4+) was done by culturing cells at a concentration of 1×106/ml in 24-well, anti-CD3 (2C11; Bio X Cell) coated plates at a concentration of 10 μg/ml followed by addition of 2 μg.ml-1 soluble anti-CD28 (37.51; Bio X Cell) with/-out 20 IU/ml of human IL-2.[2] Microparticles (5 *μ*m) or dynabeads were added to the cells at a 1:1 ratio of particles to cells and other particles(0.8 *μ*m, and 0.4 *μ*m) were added in appropriate concentration to provide the same surface area.

For flow cytometry analysis, antibodies to mouse CD4, CD8 (53-6.7), CD25 (PC61.5), CD44 (IM7), FoxP3 and CD16/CD32 (FC block) were purchased from eBioscience, BioLegend, or BD Biosciences. To study T-cell responses to various treatments T cell expansion was measured by 5-(and-6)-carboxyfluorescein diacetate, succinimidyl ester (CFSE) dilution. For CFSE dilution experiments, 5 × 105 cells were labeled with 2 μM CFSE for 13 min, washed, and plated with various particles formulations. Trypan Blue were purchased from Calbiochem. Cells were analyzed on FACSVerse using FlowJo software (Treestar).

For induced regulatory T cell (iTreg) formation experiments CD4+ T-cells were purified from mouse spleen by EasySep immunomagnetic negative selection (Stem Cell Technologies). Cells were then activated on anti-CD3e antibody (10 μg/mL) coated plates with the anti-CD28 antibody (2 μg/mL)-supplemented medium. At the same time TGF-b and IL-2 loaded particles (with different formulations) were added to the media. After four days regulatory T-cells were removed from wells coated with anti-CD3e and stained with antibodies for flow cytometry analysis. Stability of formed iTregs were also tested after 4 and 8 days by flow cytometry.

In vitro Treg Suppression assay was performed according to the previous reports. Fresh sorted iTregs were added to naïve primary CD4 T cells, which their natural Tregs (nTregs) were sorted out, at three different ratio of iTreg: naïve T cell (1:1, 1:10, and 1:30) while being stimulated with coated anti-CD3 and soluble anti-CD28 24-well plates as mentioned above. T cell cultures without iTregs were stimulated in the same manner as positive controls. Activation markers for iTregs and CD4 T cells were tested as mentioned before.

#### Statistical analysis

The presented data are expressed as average ± 95% confidence interval, unless otherwise indicated. All averages and confidence intervals are calculated by permutation methods (bootstrap).

## Supplemental Tables

**Table SI. 1.** Evaluation of SPIONs inside particles

|  | Diameter (nm) | Volume (nm<sup>3</sup>) | Relative Volume | SPION Weight Content | Average Number of SPIONs per Particles | Volume fraction of SPIONs (vol. %) | Distribution Density (nm<sup>-3</sup>) |
| --- | --- | --- | --- | --- | --- | --- | --- |
| Particle 3 | 307 | 1.5×10<sup>7</sup> | 1 | 0.25 | 5.65 | 2.43 | 3.73×10<sup>7</sup> |
| Particle 2 | 824 | 2.9×10<sup>8</sup> | 19.3 | 0.33 | 126.39 | 2.82 | 4.31×10<sup>7</sup> |
| Particle 1 | 4540 | 4.9×10<sup>10</sup> | 3234.1 | 0.36 | 22934.98 | 3.06 | 4.68×10<sup>7</sup> |

**Table SI. 2.**

|  | Diameter (nm) | Surface Area / Particle (μm<sup>2</sup>) | Total Required Surface Area (μm<sup>2</sup>) | Number of Particles to Provide 1 Particle: 1 Cell | Relative Number of Particles |
| --- | --- | --- | --- | --- | --- |
| Particle 1 | 4.5 | 63.62 | $9.5 \times 10^7$ | $1.5 \times 10^6$ | 1 |
| particle 2 | 0.8 | 2.01 | $9.5 \times 10^7$ | $4.7 \times 10^7$ | 31.64 |
| Particle 3 | 0.3 | 0.28 | $9.5 \times 10^7$ | $3.4 \times 10^8$ | 225 |
| | | | $9.5 \times 10^7$ | | |
| Dynabead | 4.5 | 63.62 | $9.5 \times 10^7$ | $1.5 \times 10^6$ | 1 |

